# Induction of proteostasis pathways in a *C. elegans* model of exercise

**DOI:** 10.1101/2025.02.21.639525

**Authors:** Lewis Randall, Gordon J. Lithgow

**Author notes:** Corresponding author: Gordon J. Lithgow.

## Abstract

Exercise is one of the most potent interventions known that is able to prevent and even treat dozens of age-related dysfunctions and diseases however much remains unknown about how its benefits are derived. Because exercise exerts such wide-ranging effects, and because a decline in protein homeostasis (proteostasis) with age has been connected to numerous age-related diseases, we hypothesized that exercise could increase the activity of proteostasis pathways like the proteasome and autophagy and that this could ameliorate age-related declines in function. We investigated the effects of exercise has on proteostasis in *Caenorhabditis elegans*. We utilized a involuntary movement exercise paradigm to investigate acute exercise and the effects of multiple days of exercise on aging and proteostasis, including the autophagy-lysosome system, the proteasome, and neurotoxic peptides. We found that exercise is able to increase proteasomal activity and autophagic flux *in vivo*, improved resistance to toxic peptides, and increased lifespan. One of the primary rationales for studying the mechanisms of exercise is to uncover potential mediators that can be repurposed to deliver the benefits of exercise to those unable to engage in physical activity. We hypothesized that exercise-elevated metabolite α-ketoglutarate, already demonstrated to improve age-related declines in flies and mice, would mimic the effects of exercise on proteostasis. Treatment with α-ketoglutarate showed resulted in complex proteostatic outcomes, indicative of the challenges in recapitulating a multi-functional domain phenomenon like exercise.

## INTRODUCTION

Exercise, the intentional engagement in physical activity, has been found to be able to delay, prevent, or treat numerous age-related diseases, including sarcopenia, diabetes, many cancers, and neurodegenerative diseases (1). The mechanisms through which these effects are achieved are complex and under increasing levels of investigation. Studies in humans have found protective benefits of exercise in a roughly dose-dependent manner, with even small amounts of activity being beneficial compared to sedentism in both middle-aged and older adults (2). However, many individuals suffering from infirmities or disabilities related (or unrelated) to age are unable to engage in sufficient exercise to experience these benefits. Mimicking the effects of exercise is desirable for these individuals, ideally with the goal of restoring them to a point where they can reengage in physical activity.

The nematode roundworm *Caenorhabditis elegans* has been a mainstay of research into aging biology due to its small size, ease of maintenance, fixed cell number, tractable genetics with a high degree of overlap with human genes, and clearly differentiable tissue types including neurons and body-wall muscle. Indeed, the musculature of *C. elegans* bears a striking resemblance in many respects to mammalian striated skeletal muscle (3). A standard procedure to measure muscle function in *C. elegans* is a so-called “thrashing assay,” where animals are placed in a small drop of isotonic buffer within which they begin thrashing involuntarily. Muscle function is determined by the number of complete side-to-side body bends the animal makes in a set time. Recently, this thrashing behavior has been characterized as equivalent in many respects to mammalian exercise (4–7). Notably, animals will thrash or “swim” for roughly 90 minutes before a refractory period, indicating exhaustion of the animals, before swimming resumes (4). Exposing *C. elegans* to repeated bouts of swimming has been shown to elicit adaptations akin to those in mammals, including increased maintenance of mitochondrial networks and function, increased chemotaxis (a test of neuronal health), resistance to certain proteotoxic stresses, and increased median lifespan (4,5). Research using this model has also established that the effects are not due solely to a lack of food while swimming (no bacterial food source is present in the swimming media), with benefits shown beyond that of food restriction as well as while swimming in the presence of food (6,7). So-called “endurance” exercise in mammals is characterized by usage of lipid catabolism for generation of ATP, as opposed to “resistance” exercise which is characterized by glycolysis (8). In the *C. elegans* thrashing exercise paradigm, quantitative PCR of genes encoding metabolic functions found an elevation in abundance of mRNAs related to lipolysis and fatty acid oxidation and a decrease in mRNA abundance of proteins involved in glucose transport and glycolysis (5), indicating a similarity to “endurance” exercise in mammals.

A key feature underlying many aging conditions is the accumulation of damaged, aggregating proteins (9–13). While this is most well-characterized in neurodegenerative diseases like Alzheimer’s (14,15), the accumulation of insoluble protein has been found to be a general feature of aging across multiple organs and model organisms (9,16). This loss of proteostasis with aging is accompanied by a decline in function of cellular recycling mechanisms including autophagy and the proteasome and activation of these mechanisms has been found to delay aging and reduce the development of age-related morbidities (17–22). Intriguingly, exercise has been found to induce autophagy in humans as well as in rodent models of exercise, including tissues not directly involved in exercise like the brain (23–25). Less is known about the effect of exercise on the proteasome, but a study in humans found that exercise attenuated hyperactivation of the proteasome in patients with muscle dystrophies (26) while a study in mice found that exercise increased hippocampal proteasome activity (19). More broadly, exercise and its potentially mitigating role in the decline of proteostasis with aging remains a surprisingly understudied area.

Exercise is metabolically intensive and the breakdown of energy sources during ATP production, primarily lipids and carbohydrates, generates numerous byproducts. Far from being waste products, these have important functions in metabolic pathways and as signaling molecules. Several studies have analyzed the acute metabolome of exercise in humans and rodent models in search of so-called “exerkines” – molecules elevated in response to exercise – that could potentially be used to replicate the effects of exercise (27–29). One molecule that shows consistent elevation after exercise is the TCA cycle intermediate alpha-ketoglutarate (AKG). Intriguingly, AKG has already been shown in *C. elegans, Drosophila*, and mice to extend lifespan along with ameliorating a number of age-related dysfunctions and the compression of age-related morbidities (30–32).

In this study, the *C. elegans* swimming exercise model was used to determine what effect exercise has on different measures of proteostasis, including autophagy, the proteasome and the reduction of known neurotoxic peptides with both acute (1-hour) and chronic exercise. Notably, supplementation with the TCA cycle intermediate AKG, a known exerkine, mimicked some, but not all, of exercise’s effects on proteostasis. Lastly, we explore a potential role for the regulation of protein acetylation by the metabolic enzyme ACS-19 and transcription factor HLH-30 in mediating the effects of exercise on proteostasis.

## RESULTS

### Thrashing exercise in *C. elegans* resembles mammalian endurance exercise

Exercise in humans is typically categorized by intensity and duration on a rough scale from resistance (relying primarily on glycolysis) to endurance (relying primarily on fatty acid oxidation) (8). Because prior *C. elegans* exercise studies found an upregulation of lipolysis and fatty acid oxidation genes after a bout of swimming and a downregulation of glycolytic genes, we hypothesized that there would be a decrease in lipid stores. Using the exercise modality depicted in **Figure 1A** followed by lipid staining with Nile Red, exercised animals showed a reduction in fat stores (**Figure 1B**). Surprisingly, animals exposed to food-restriction showed no decrease in fat stores relative to fed animals. Unlike mammals, *C. elegans* do not have specific fat storage tissues, instead storing fats in the form of lipid droplets within cells, primarily the intestine, muscle, and hypodermis. Proteins involved in the packaging and breakdown decorate the surface of lipid droplets. The oxidoreductase DHS-3, ortholog of human SDR16C5, is an intestine-specific lipid droplet protein; using a GFP-tagged endogenously expressed version, intestinal lipid droplet content was measured and found to be decreased in a single group of animals before and after a single 60-minute bout of exercise (**Figure 1C**). The Perilipin ortholog, PLIN-1, is involved in packaging and maintaining lipid droplets in the intestine, body-wall muscle, and hypodermis. By using an mCherry-tagged version of the protein, fluorescence intensity of the construct was used to measure lipid droplet content before and after a 60-minute bout of exercise (**Figure 1D**). Unlike a prior study using the *C. elegans* exercise model (6), we found that fat stores measured by the intensity of lipid droplet fluorescence decreased primarily in the intestine (**Figure 1C**) after an acute bout of exercise as opposed to the muscle (5), where there was a small but insignificant decrease.

**Figure 1.**
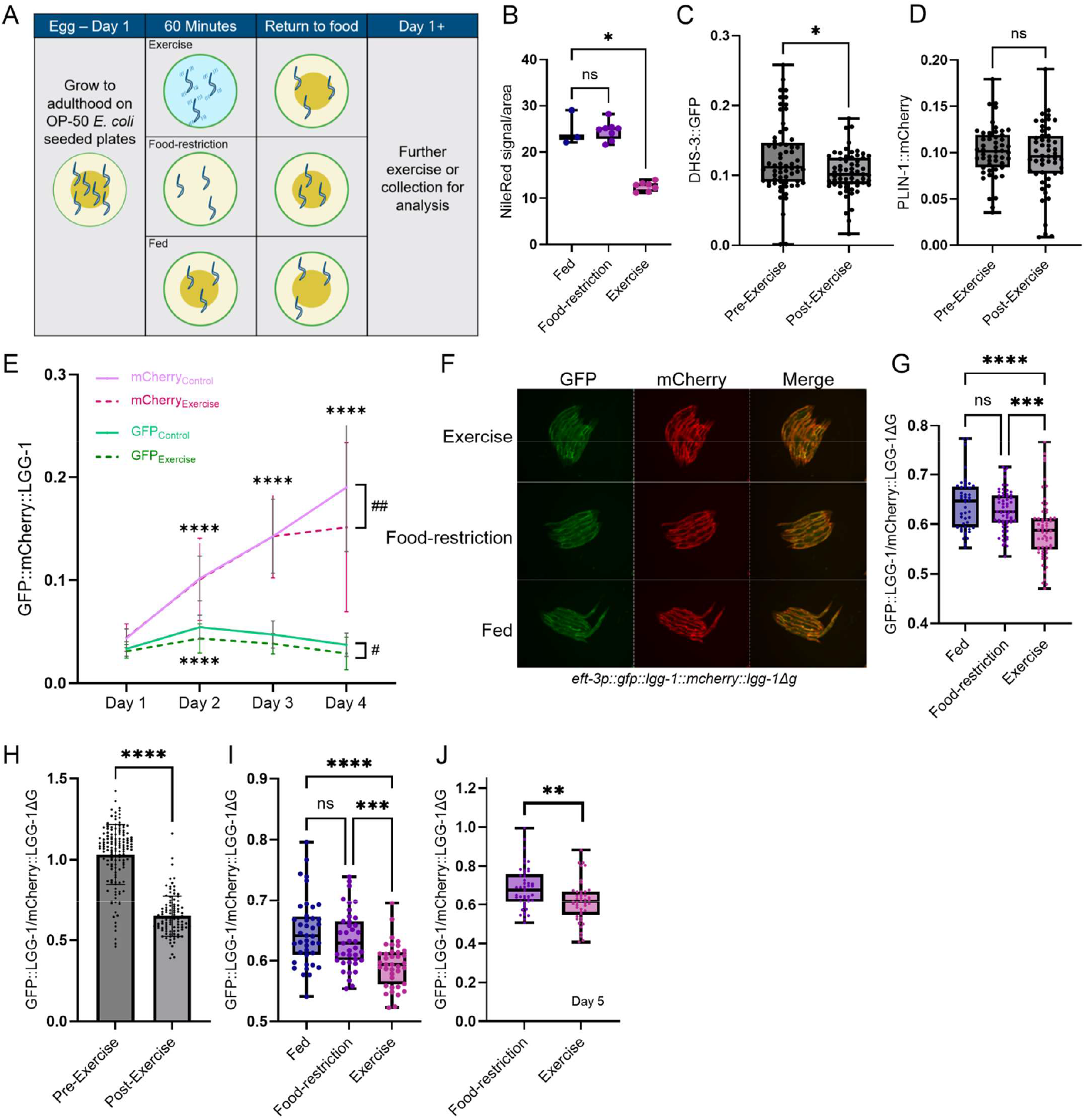
**(A)** Schematic of *C. elegans* thrashing exercise paradigm. **(B)** Nile Red fluorescence intensity of animals exposed to Fed, Food-restriction, or Exercise conditions for 1 hour (n=4-6) and analyzed by one-way ANOVA. Measurement of lipid droplet fluorescence in the intestine **(B)** and intestine, hypodermis, muscle and embryo **(C)** in a single group of animals before and after 1 hour of exercise. **(E)** Signal from a GFP::mCherry::LGG-1 whole body-expressed construct from Day 1 through Day 4 of adulthood in Control (solid line) and Exercise (dashed line) groups. **(F-J)** Representative images of *C. elegans* with an LGG-1::GFP::LGG-1ΔG::mCherry endogenously-expressed construct exposed to Fed, Food-restriction, or Exercise condition for 1 hour with quantification of the GFP/mCherry ratio in **(G)**, quantification in a single group of animals before and after 1 hour of exercise **(H)**, quantification in Day 2 animals after 2x Fed/Food-restriction/Exercise on Day 1 **(I)**, and quantification in animals exposed to Food-restriction or Exercise on Day 5 only **(J)**.

### Exercise in *C. elegans* induces autophagy

Macroautophagy, hereafter autophagy, is the primary cellular recycling mechanism via which damaged organelles, proteins, and other macromolecules are broken down for reuse. A decline in autophagy with age has been tied to multiple age-related diseases and its activation has been found to be required for the lifespan-extending effects of multiple interventions (17,21). Exercise has been shown to induce autophagy in rodent models of exercise as well as in humans through tracking changes to the abundance of the mature (lipidated) and immature forms of the autophagic membrane protein LC3 (*C. elegans* ortholog LGG-1) (21,24,33–36), but an increase in autophagic flux after acute exercise has yet to be demonstrated. Because of the pulsatile nature of the effects of exercise (28,37), there may be acute responses to exercise that are important in addition to adaptations to chronic exercise; a burst of autophagy to clear damaged macromolecules may be one of these acute events.

*C. elegans* expressing *gfp::mcherry::lgg-*1 have been used to track autophagic function with age via analysis of autophagosomes and autolysosomes. Autophagosomes appear as yellow puncta from combined GFP and mCherry fluorescence and turn to red puncta as they fuse with acidic lysosomes and the pH-sensitive GFP signal is quenched. By using confocal microscopy to count autophagosomes (yellow puncta) and autolysosomes (red puncta) in different *C. elegans* tissues, Chang *et al* found that in most tissues there is an age-related accumulation of dysfunctional autolysosomes (38). To determine whether this age-related decline in autophagy could be observed *in vivo*, we used a laser-based bulk object sorter (COPAS Biosorter™) (39) to track the same group of animals over time. Starting from Day 1 of adulthood, fed animals and animals exposed to 1 hour of exercise per day were measured daily until Day 4 of adulthood (**Figure 1E**). Interestingly, we observed an age-related increase of red fluorescence but an increase of green fluorescence only on Day 2, which closely matches the pattern of puncta-based analysis of autophagosome and autolysosome accumulation, respectively (38). In animals exposed to 1 hour of exercise per day, there was a small but significant decrease in the accumulation of red fluorescence (potential dysfunctional autolysosomes) and of green fluorescence (potential autophagosomes) (**Figure 1E**).

While the above experiment demonstrated an age-related decline in autophagy and a slight amelioration with 1 hour of exercise per day in *C. elegans*, whether exercise is able to acutely induce autophagic flux *in vivo* has yet to be determined. Autophagic flux *in vivo* in whole animals can be determined with the *lgg-1*:*gfp*::*lgg-1*ΔG::mcherry construct (40,41). When activated, the enzyme ATG-4.1 cleaves the C-terminal glycine of the wild-type LGG-1, generating equimolar pools of GFP::LGG-1 and mCherry:: LGG-1ΔG (40,41). The mutant LGG-1ΔG lacks this C-terminal glycine, preventing ATG-4.1 activity, subsequent lipidation, and incorporation into the autophagosome membrane (40). By fusing GFP to the normal LGG-1 and an mCherry to the mutant version, the ratio of green-to-red signal can be used as a proxy for autophagic activity as the green signal is quenched within the acidic autolysosome and the red signal remains unchanged in the cytosol. we subjected this strain, EQ1301, to the swimming exercise protocol along with fed and food-restriction controls to observe the acute effect of exercise on autophagy. Indeed, one hour of swim exercise increased autophagic flux relative to both the fed group as well as the food-restriction group (Images in **Figure 1F**, quantification in **Figure 1G)**. Surprisingly, one hour of food-restriction had no effect on the level of autophagic flux.

The COPAS flow sorter system provides for high-throughput analysis of large objects, including live *C. elegans*, expressing or marked with fluorescent tags. The COPAS provides for recovery of live animals, allowing them to be reanalyzed. In a follow-up experiment using the EQ1301 strain, Day 2 adult animals were analyzed via COPAS, measuring whole-body green and red signals, and recovered in S-basal, maintaining the worms in a swimming exercise condition. After one hour of swimming exercise, the same population of worms was reanalyzed via the COPAS. The GFP::mCherry ratio showed a significant decrease after 1 hour of exercise (**Figure 1H**). This orthogonal *in vivo* approach showed an acute increase in autophagic flux within the same population of animals.

In addition to the acute effects of exercise on autophagy, it was worthwhile to determine whether swimming exercise in *C. elegans* can increase the basal level of flux 24 hours post-exercise to better match studies done in mammalian systems. To test this, Day 1 adult *C. elegans* were subjected to two bouts of 1 hour swimming exercise, food-restriction control, or fed conditions and then collected for imaging ~24 hours after the second bout on Day 2 of adulthood for imaging. Similar to mammalian chronic exercise studies, only the exercised animals had an increased level of basal autophagic flux (**Figure 1I**). This indicates that chronic exercise in *C. elegans*, similar to what has been observed in rodent exercise, has enduring effects on autophagy. Lastly, to see whether older animals were still capable of exercise-induced autophagy, Day 5 adult animals were exposed to 1 hour of exercise or equivalent food-restriction and exercise likewise increased autophagic flux above that of food-restriction (**Figure 1J**).

### Exercise in *C. elegans* improves proteasomal function

Degradation of damaged proteins as well as baseline protein turnover by the proteasome is an additional mechanism through which cells maintain proteostasis. Cargo is tagged for degradation by the proteasome via E3-ligase-mediated ubiquitination to create polyubiquitin chains, with repeated K48 ubiquitination as the primary signal for degradation by the proteasome (20). To quantify proteasomal function *in vivo* in *C. elegans*, we used a strain (GR3090) expressing a GFP constitutively tagged with a non-cleavable ubiquitin (GFP::Ub) in body-wall muscle. Under normal conditions, GFP::Ub is produced and sent for degradation via the proteasome. Upon proteasomal dysregulation, GFP::Ub accumulates within cells, leading to an increase in GFP fluorescence. Because the proteasome is known to become dysfunctional with age (20,44), we first tested whether chronic exercise in *C. elegans* was able to counteract it. Animals were exercised 2 times per day for Day 1 through Day 4 of adulthood and were collected for quantification of GFP::Ub fluorescence on Day 5 and Day 7 of adulthood, 1 and 3 days post-exercise, respectively (**Figure 2A**). Exercise significantly reduced the accumulation of GFP::Ub fluorescence relative to food-restriction control animals. Although GFP::Ub accumulation continued from Day 5 to Day 7 even in exercised animals, it was significantly lower than non-exercised Day 7 animals, even 3 days post-exercise. Additionally, another group of animals was fed only from Day 1 through Day 6 and then exposed to 1 hour of exercise or food-restriction on Day 7. Exercise significantly reduced the GFP::Ub signal relative to food-restriction, to the same level as the animals with 4 days of exercise (**Figure 2A**).

**Figure 2.**
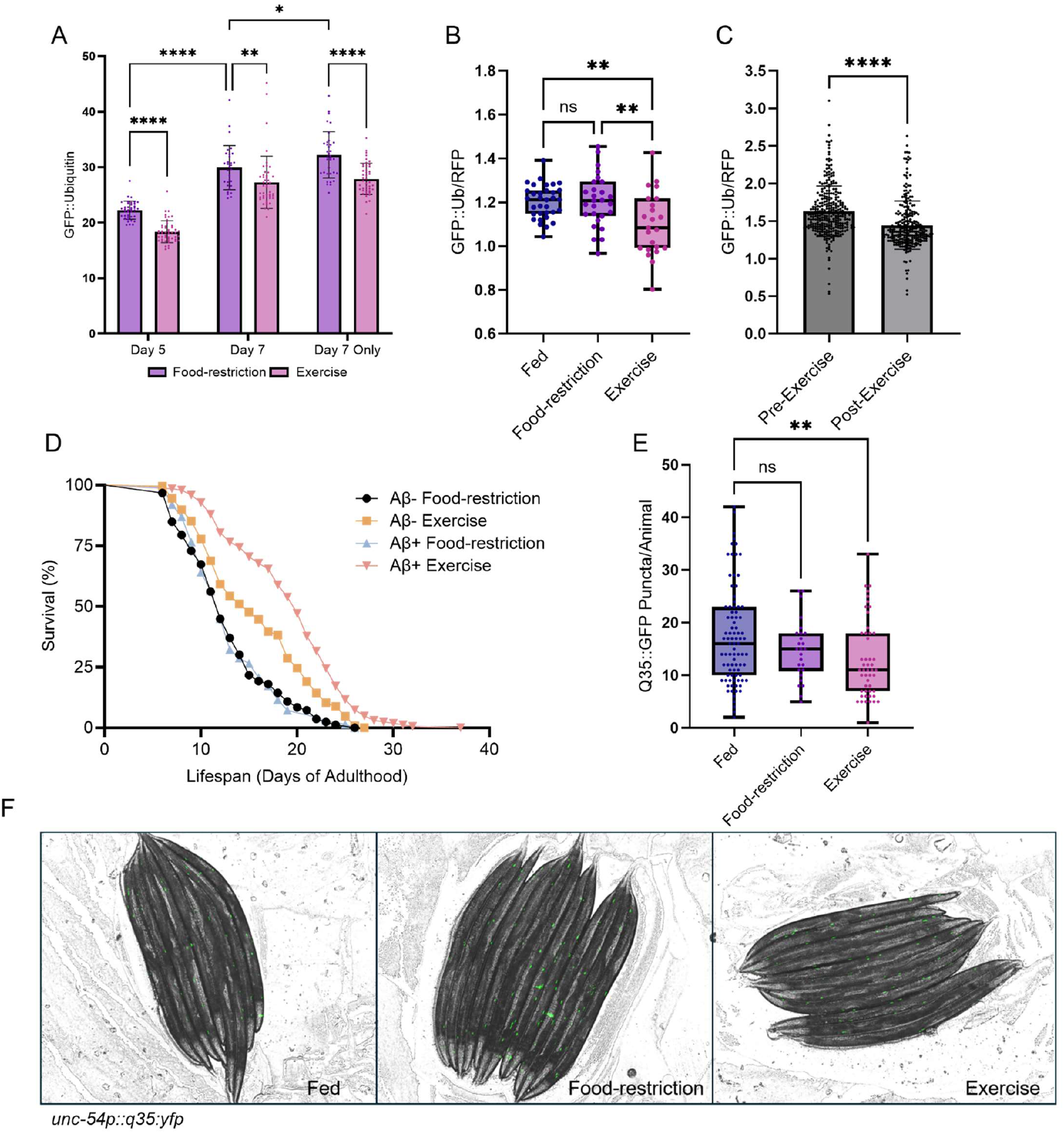
**(A)** Quantification of GFP/RFP signal in *C. elegans* expressing non-cleavable Ubiquitin tagged to GFP and non-tagged RFP in muscle in Day 1 animals after exposure to Fed, Food-restriction, or Exercise for 1 hour. **(B)** Quantification of the GFP/mCherry ratio in the same strain as **(A)** in a single group of Day 1 animals before and after 1 hour of exercise. **(C)** *C. elegans* expressing only the GFP::Ubiquitin construct were treated with Food-restriction or Exercise 2x/Day from Day 1 through Day 4, then quantified on Day 5 and Day 7, with an additional group only Fed from Day 1 through Day 6 and treated with Food-restriction or Exercise 2x on Day 7 only. **(D)** Lifespan analysis of *C. elegans* expressing and not expressing pan-neuronal human Aβ_1-42_ that were Food-restricted or Exercised 2x/Day from Day 1 through Day 5 (3 biological repeats, n=89-186). **(E)** Quantification of the number of Q35::YFP puncta per worm in **(F)**. Representative images of Day 4 *C. elegans* expressing Q35::YFP polyglutamine tracts in muscle after 3 Days of 2x/Day Fed, Food-restriction, or Exercise condition.

I then used a strain (PP607) that expressed the same GFP::Ub construct along with an RFP which is expressed via the same promoter as the GFP::Ub construct (*unc-54*). The RFP fluorescence can then be used as an internal control, with a lower GFP::RFP ratio indicating higher proteasomal activity. In this strain, exercise on Day 2 of adulthood decreased the GFP::RFP ratio relative to food-restriction and fed animals, indicating an acute increase in proteasomal activity with exercise (**Figure 2B**). Additionally, a single bout of exercise was found to reduce the GFP::Ub::RFP ratio in a single group of animals before and after 1 hour of exercise (**Figure 2C**), again indicating that exercise is able to acutely increase proteasomal activity.

### Exercise in *C. elegans* modulates cytotoxic proteins

A striking commonality in the etiology of many age-related diseases is the accumulation of misfolded, damaged proteins. In *C. elegans* expression of human neurotoxic proteins often leads to impairments in movement, chemosensation, and lifespan (12,45–47). Epidemiological studies have found a substantial reduction in the risk of developing Alzheimer’s Disease (AD) among frequent exercisers versus their more sedentary counterparts. As a measure of the ability to handle this toxic peptide, we decided to test whether extended exercise could increase lifespan in a *C. elegans* strain expressing human amyloid-β_1-42_ pan-neuronally (GRU102) (48). This strain and their non-amyloid matched strain (GRU101) were exercised 3x/day from Day 1 through Day 5 of adulthood or food-restricted for the same periods. After the last bout of exercise, animals were moved to fresh plates and tracked for lifespan (**Figure 2D**). In the non-amyloid-β_1-42_ expressing control animals, exercise significantly increased median lifespan from 12 to 15 days (a 25% increase) while maximal lifespan was unchanged. Unexpectedly, exercise of the amyloid-β_1-42_ expressing animals significantly extended median lifespan beyond that of non-amyloid-β_1-42_ exercised animals from 12 to 20 days (a 67% increase) as well as extended maximal lifespan from 26 to 37 days (a 42% increase). This study was done 3 separate times with the same pattern appearing in each biological repeat. Intriguingly, prior work with this strain has found that treatment with lithium also increases lifespan of the amyloid-β_1-42_ animals beyond that of the non-amyloid-β_1-42_ controls (48). The exact mechanism for this remains unknown. However, the level of expression of amyloid-β_1-42_ in this strain is somewhat low and, when given a proteostasis-enhancing treatment such as exercise, may potentially act as a mild hormetic stress.

The Huntingtin protein is known to aggregate and cause Huntington’s Disease in humans when a CAG motif coding for glutamine expands beyond ~35 repeats. The polyglutamine tract alone will aggregate and cause toxicity when expressed in *C. elegans* body-wall muscle under control of the *unc-54* promoter (45). The *C. elegans* exercise regimen was tested for its ability to affect the aggregation of this toxic peptide using animals expressing Q35::YFP in body-wall muscle. In Day 1 animals, 2 1-hour bouts of exercise significantly lowered the number of Q35::YFP puncta per worm relative to animals exposed to food-restriction or left as fed (**Figure 2E**).

### Supplementation with the exercise-induced metabolite alpha-ketoglutarate mimics effects on autophagy but not the proteasome

Metabolomic studies of both human and rodent model exercise have identified numerous metabolites both up- and down-regulated after a bout of exercise. One group of metabolites that is consistently elevated after exercise are TCA cycle intermediates. Among those, we and others found that alpha-ketoglutarate (AKG) ameliorated age-related morbidities and extended lifespan in *C. elegans, Drosophila*, and mice (30–32). Here, supplementation with AKG was tested in *C. elegans* to determine its effects on proteostasis mechanisms as well as handling of toxic peptides. Using the LGG-1-GFP::LGG-1ΔG::mCherry reporter strain, Day 1 adult animals were placed on plates containing 8 µMAKG or control plates. When collected and imaged 24 hours later, AKG-treated animals showed higher autophagic flux than controls (**Figure 3A**). Surprisingly, AKG-treated animals had decreased proteasomal function, shown via increased accumulation of GFP::Ub (**Figure 3B**). Additionally, AKG-treated animals had a higher number of polyglutamine puncta than control animals (**Figure 3C**), indicating reduced ability to handle this toxic peptide (45). Despite worsened function in two out of three measures of proteostasis, AKG-treated animals showed an extension in median lifespan (**Figure 3E**).

**Figure 3.**
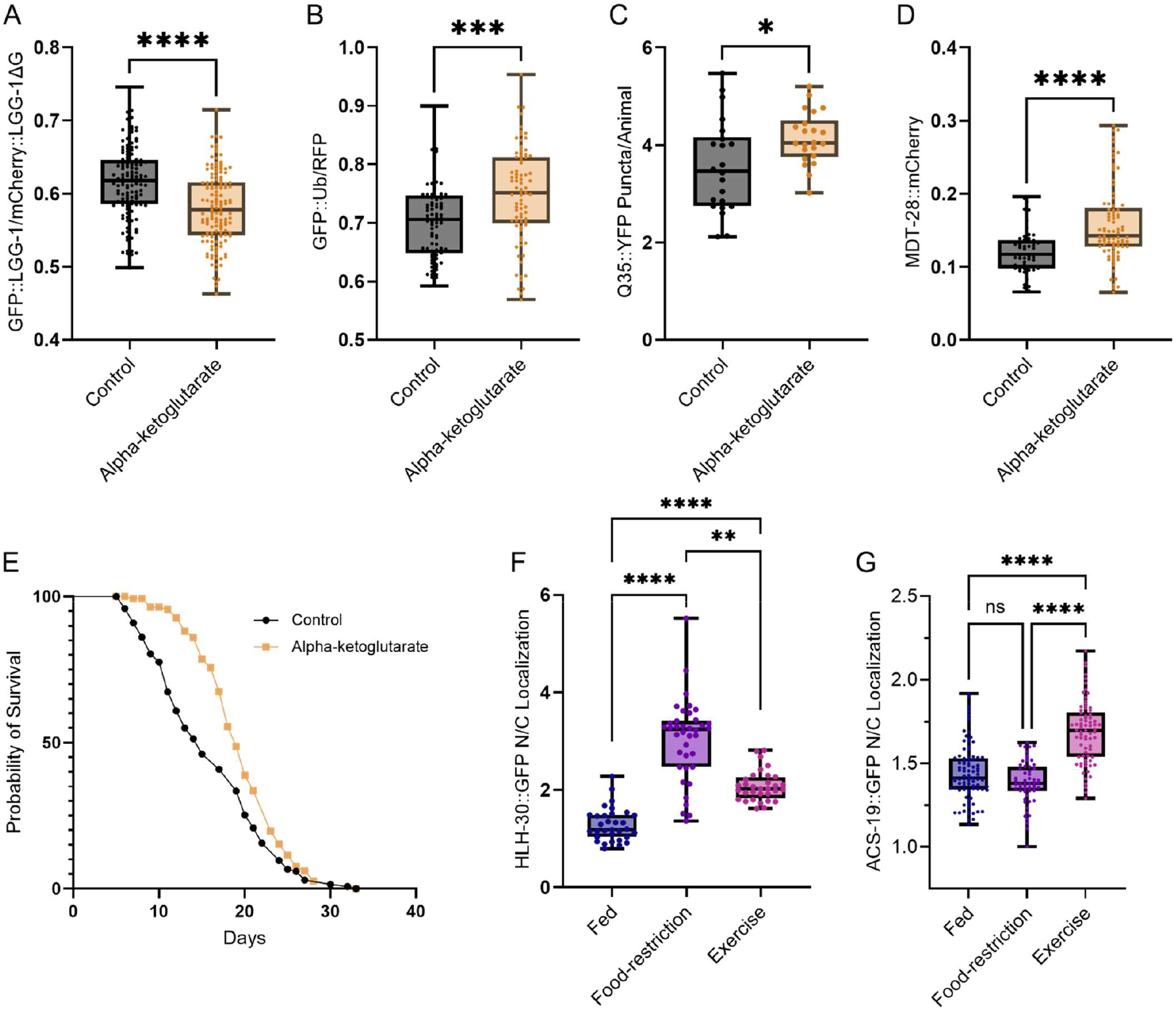
**(A)** Quantification of the GFP/mCherry ratio in *C. elegans* expressing the LGG-1::GFP::LGG-1ΔG::mCherry construct placed on 8µMAKG or Control plates on Day 1 and imaged on Day 2. **(B)** Quantification of the GFP/RFP ratio in *C. elegans* expressing non-cleavable Ubiquitin tagged to GFP and non-tagged RFP in muscle placed on 8µMAKG or Control plates on Day 1 and imaged on Day 2. **(C)** Counts of polyglutamine aggregate in body wall muscle (Q35::YFP) per worm in *C. elegans* grown on plates containing 8µMAKG or Control from Day 1 through Day 3 and imaged on Day 4. **(D)** Lipid droplet fluorescent intensity in intestine, hypodermis, muscle, and embryos after treatment with 8µMAKG or Control from Day 1 to Day 2. **(E)** Lifespan analysis of N2 *C. elegans* on plates containing 8µMAKG or Control starting from Day 1. **(F)** Quantification of nuclear versus cytosolic localization of HLH-30::GFP after 1 hour of Fed, Food-restriction, or Exercise condition. **(G)** Quantification of nuclear versus cytosolic localization of ACS-19::GFP after 1 hour of Fed, Food-restriction, or Exercise condition.

### Nuclear localization of autophagy-regulating proteins after exercise

The master regulator of autophagy, transcription factor TFEB/HLH-30, translocates to the nucleus where it drives transcription of components of the autophagy/lysosome system. In *C. elegans*, the TFEB orthologue, HLH-30, is known to quickly translocate to the nucleus after a mild stress, such as removal from food. Indeed, as expected, an endogenously tagged HLH-30::GFP showed a high degree of nuclear localization in intestinal cells after both exercise and food-restriction, in contrast to fed animals (**Figure 3F**). Despite the localization of HLH-30 to the nucleus under food-restriction conditions, one hour was insufficient to increase autophagic flux (**Figure 1G**). This raised the possibility that exercise induces the recruitment of additional proteins involved in the regulation of autophagy. One such enzyme is ACS-19, the *C. elegans* ortholog of ACSS2, which is responsible for generating acetyl-CoA from acetate in the cytosol and nucleus. Upon localization to the nucleus, it produces acetyl-CoA at the sites of histone acetylation. Notably, ACSS2 is known to co-localize with TFEB in the nucleus to positively regulate the expression of autophagy and lysosomal genes (49). In the study by Li et al, they found that upon glucose deprivation in cell culture, AMPK phosphorylated ACSS2, leading to its nuclear translocation and binding to TFEB where they bound to the promoter regions of lysosomal and autophagy genes. In in the cytoplasm, it has been connected to acetylations of autophagic proteins that promote their activity, including SQSTM1/p62(50). we decided to test, therefore, whether exercise would lead to ACS-19 nuclear translocation to help explain the observed increase in autophagic flux with exercise. Using a GFP-tagged ACS-19, localization was shown to increase in the nucleus of intestinal cells after acute exercise relative to food-restriction and fed controls (**Figure 3G**), consistent with what was seen in human cell culture under glucose deprivation, although further validation is required to show binding with HLH-30 following exercise.

## METHODS

### C. elegans strains and maintenance

*C. elegans* were grown following standard maintenance protocols. In brief, animals were fed OP-50 strain of *E. coli* seeded onto NGM agar plates. OP-50 *E. coli* was cultured in Miller LB Broth (EMD) overnight at 37°C. NGM agar was made by autoclaving Difco Agar (BD) with NaCl (Sigma-Aldrich), Bacto Peptone (BD), and milliQ H_2_O. After cooling to ~55°C, KPO_4_ (Sigma-Aldrich), MgCO_4_ (Sigma-Aldrich), CaCl_2_ (Sigma-Aldrich), and cholesterol (Sigma-Aldrich) were added and plates were poured. After drying at room temperature, plates were seeded with the OP-50 *E. coli*. Animals were synchronized via egg lay of Day 1 or Day 2 gravid adults for <2 hours. Strains used for experiments were Bristol N2 (wild-type); AM140 (*unc-54p::q35::yfp*); BC10604 (*acs-19p::acs-19::gfp*); EQ1301 (*lgg-1p::lgg-1::gfp::lgg-1*Δ*g::mcherry*); GMC101 (*unc-54p::abeta*_*1-42*_); GR3090 (*unc-54p::ubiquitin::gfp*); GRU101 (*myo-2p::yfp*); GRU102 (*myo-2p::yfp*; *unc-119p::abeta*_*1-42*_); LIU1 (*dhs-3p::dhs-3::gfp*; *unc-76*); LIU2 (*mdt-28p::mdt-28::mcherry*; *unc-76*); MAH349 (*sqst-1p::sqst-1::gfp + unc-122p::rfp*); MAH704 (*rgef-1p::sqst-1::gfp + unc-122p::rfp*); MAH215 (*lgg-1p::mcherry::gfp::lgg-1*; *rol-6*); MAH235 (*hlh-30p::hlh-30::gfp*; *rol-6(su1006)*); PP607 (*unc-54p::ubiquitin::gfp; unc-54p::rfp*). EQ1301, MAH215, MAH349, and MAH704 were kind gifts from the Hansen Lab. All other strains were obtained from the *Caenorhabditis elegans* Genetics Center (CGC) at the University of Minnesota(51).

### C. elegans exercise

This *C. elegans* swimming exercise protocol has been adapted from that developed by the Driscoll laboratory(4,5) and the Andersen laboratory(52). A part of standard *C. elegans* handling is the use of liquid transfers, where an isotonic solution such as M9 or S-basal buffer is used to transport many animals from plate to plate. *C. elegans* are sensitive to osmotic stress and the use of pure water will result in the rapid death of the animal. When *C. elegans* are placed in an isotonic solution they begin an involuntary thrashing consisting of bending the body in one direction and then the other. Measuring this thrashing rate (the number of complete body bends in a set period of time) is used as a common proxy for overall animal health. S-basal buffer was used for all exercise experiments and was made by combining 20mL 5M NaCl, 50mL 1M phosphate buffer, and 930mL H_2_O and autoclaving.

To perform the exercise regimen, worms are picked from a synchronized population onto an NGM agar plate that has not been seeded with bacteria, taking care to minimize the amount of bacteria transferred. Once the worms have been transferred, the plate is flooded with S-basal buffer, with the volume corresponding to the size of the plate used (1mL for a 3cm plate, 2mL for a 6cm plate, 5mL for a 10cm plate). For the food-restriction control, worms are transferred to an unseeded NGM plate the same as with the exercise group, but only a small drop of S-basal is put on the worms to help wash off any bacteria that has been transferred along with the worms. For the fed control group, worms are transferred to a normal, seeded NGM plate. All plates sit on the bench facing upright, with their lids on, covered with a sheet of aluminum foil to protect them from light (exposure to light has been found to reduce *C. elegans* lifespan(53)). The duration for each bout of exercise is one hour unless stated otherwise. At the end of the hour, the exercise group is collected via a glass pipettor and moved to a 1.5 mL Eppendorf tube where they are allowed to settle; excess S-basal is removed and the remaining liquid with the *C. elegans* is transferred to a fresh, seeded, NGM plate. The food-restriction control group is moved to a fresh, seeded, NGM plate via picking with bacteria as normal. For some experiments, animals were exercised multiple times in a day, and in these cases, animals were allowed to rest for 2-3 hours between bouts. Because *C. elegans* become increasingly frail with age and more care needs to be taken moving them, animals were typically not exercised past Day 5 of adulthood. To avoid interference with development and associated pathways, animals were not exercised prior to Day 1 of adulthood.

For experiments using the COPAS™ object sorter, animals were collected by flooding the plate with S-basal, transferred to a 50mL conical tube, and measured by passing through the detection chamber. Then animals were dispensed into plate wells filled with S-basal so that the animals continued thrashing. After 1 hour of exercise in the wells, animals were transferred via pipettor to a 50mL conical and re-analyzed by the COPAS™. Depending on the experiment, animals were either disposed of or, for multi-day experiments, dispensed onto seeded NGM plates.

### Lifespan assays

Lifespan assays were performed by moving animals daily to new plates from Day 1 through Day 5 of adulthood, avoiding carryover of eggs and larvae, at which point scoring for survival began(54). If food became depleted, animals were moved to fresh plates. Any larvae that hatched post-Day 5 were removed. Plates were checked daily, and death was ascertained via gentle taps with a platinum wire pick to the body and then head of immobile animals. Animals with internal hatching (“bagging”) were censored. No FuDR to prevent the hatching of newly laid eggs was used in any lifespan experiment.

### Imaging and quantification

For whole-animal imaging, animals were placed in droplets of 2.5mM levamisole or 20mM sodium azide dissolved in M9 buffer on 2% agar pads on glass slides. Once paralyzed, animals were aligned on the pad using a platinum wire or eyelash. Imaging was performed with an AX10 Zeiss microscope and Excelitas X-Cite 120 LED lamp at 100% power. For nuclear localization experiments, animals were placed into slide wells containing 2.5mM levamisole with a coverslip laid atop to bring the intestine into a single focal plane. Quantification was performed using Fiji (ImageJ). For whole-animal quantification, fluorescence intensity was determined by using intDen normalized to body size. For nuclear-cytoplasmic localization, two spaces within the cytoplasm and one within the nucleus were measured, the cytoplasmic measurements averaged together and the ratio of nuclear to cytoplasmic determined by dividing the nuclear signal by the cytoplasmic signal.

Nile Red staining was done following the protocol described in Stuhr et al(55). In brief, animals were collected in PBST (PBS + 0.01% Triton X-100 (Sigma-Aldrich)) and rinsed to remove bacteria. Animals were then fixed in 40% isopropanol for 3 minutes. Nile Red stock solution was added to 40% isopropanol (6µLof 0.5 mg/mL into 1mL isopropanol). Animals were then incubated in the Nile Red solution for 2 hours in the dark followed by a rinse and 30 minute incubation with PBST. Animals were allowed to settle by gravity (~30 seconds), the supernatant was removed, and animals were placed on a microscope slide for imaging in the GFP/FITC channel.

### Worm sorter (COPAS™)

In addition to the plate-based exercise method described above, *C. elegans* exercise was performed in tandem with the (Complex Object Parametric Analyzer and Sorter) system from Union Biometrica worm-sorter machine. The COPAS™ is a flow cytometry-like system for analyzing objects too large for conventional cell analyzers and sorters. It allows for high-throughput analysis of organoids, *Drosophila* larvae, *C. elegans*, and other similarly sized objects. Like cell sorters, it allows for the safe recovery of objects passed through, including live *C. elegans*. Laser power was determined for each fluorophore per strain prior to each experiment (remaining consistent between biological repeats). To analyze a single population of animals before and after a single one-hour bout of exercise, age-synchronized *C. elegans* were collected in S-basal by flooding the plate and picking them up with a glass pipettor and moved to a 50mL Eppendorf tube. S-basal buffer was then added to the tube to bring the volume to >5mL and attached to the machine for analysis. Quantification was performed by normalizing the signal for the respective fluorophore channel by the time-of-flight (TOF) for each event, a proxy for body size.

### Paralysis assay

The GMC101 strain (*unc-54p::*_*aβ1-42*_::*unc-54 3’-UTR* + *mtl-2p::gfp*) is widely used amongst *C. elegans* researchers studying proteostasis or Alzheimer’s Disease. The strain expresses human amyloid-β_1-42_ under control of the muscle *unc-54* transcriptional promoter. When moved from incubation at 20°C to 25°C on Day 1 of adulthood, the animals paralyze within 24-36 hours as a result of amyloid-β aggregation(46). For these experiments, L4 stage animals were placed on NGM plates containing 20µM NU9056. On Day 1 of adulthood, the plates were temperature-shifted to 25°C. On Day 2 of adulthood, the plates were scored when at least one condition appeared to show 60-80% paralysis. Once at least one condition showed that degree of paralysis, all conditions were scored. To score plates, animals that did not move upon gentle touch with a platinum wire were placed on one side of a plate. If an animal had not moved from its new position when re-checked 5 minutes later the animal was counted as paralyzed.

### Statistical analyses

All statistical analyses were carried out in GraphPad Prism™ 10 (GraphPad Software, LLC). Two-tailed Student’s *t*-tests were used for 2-group comparisons unless standard deviations between groups were significantly different, in which case a Welch’s correction was included. If the distributions did not follow a Gaussian distribution a two-tailed, non-parametric Mann-Whitney test was used. For comparisons of 3 or more groups, one-way ANOVA, with Welch’s correction for significantly different standard deviations and a Kruskal-Wallis non-parametric test if the data did not follow a Gaussian distribution, followed by the appropriate multiple comparisons test (Tukeys, Dunnett’s T3, or Dunn’s).

## DISCUSSION

Exercise studies in *C. elegans* represent a small but growing field(4–6,52,56–61), with exercise paradigms including swimming and different vibration-induced thrashing methods all showing improvements in function at both cellular and physiological levels. The ability of these exercise paradigms to replicate many aspects of mammalian exercise is striking, including maintenance of mitochondrial networks(5,7,58), improvements in age-related muscle function decline(5,62), and resistance to proteotoxic stress(5,57,58). Additionally, one study suggested that, akin to mammalian exercise, AMPK (AAK in *C. elegans*) regulates some aspects of the response to exercise(7) and like in mammals, constitutive activation of AMPK/AAK recapitulates certain benefits of exercise pertaining to mitochondrial health. To add to this growing body of work, we decided to focus on exercise and its impact on age-related declines in proteostasis. While prior studies of *C. elegans* exercise have observed reductions in toxicity from amyloid-β and polyglutamine repeats, they used distinct models of these toxic peptides and tested different impacts on organismal health than those tested here. First, we looked at lipid stores after exercise and confirmed their use during exercise, both as whole-body fat stores with Nile Red staining and with tissue-specific lipid droplets. In the lipid droplet experiments, it appears that, contrary to prior studies of *C. elegans* exercise, the intestinal lipids may be used more than lipids stored in the body-wall muscle. Interestingly, recent research has found an insulin-like protein in *C. elegans* that blocks the breakdown of fat stores during periods of food deprivation. The protein, INS-7, is secreted by intestinal cells upon fasting and travels to ASI neurons where it reduces the release of a neuropeptide that drives fat breakdown in peripheral tissues(63). This study is consistent with the results described above, where short-term food-restriction did not result in measurable fat loss. Because of the involuntary nature of the thrashing behavior in *C. elegans*, it must be able to override this INS-7 signaling in order to make fat stores available for use. An interesting follow-up experiment could test the fluorescently labelled INS-7::mCherry strain used in their study under the exercise and food-restriction paradigm(63). The presence or absence of secretion of INS-7::mCherry by the intestine and its migration to neurons in response to exercise will help to inform in which tissue this signaling pathway is being blocked. If there is no INS-7::mCherry, it may be being blocked in the intestine, while if there is INS-7::mCherry in the neurons, then the neurons may have other peripheral signals coming in to increase production and release of the neuropeptide driving fat breakdown.

Additionally, through these experiments, we showed that the *C. elegans* exercise paradigm is able to ameliorate age-related loss of proteostasis in multiple systems. On one end, the proteosome and autophagy-lysosome systems were assayed, and their activities were shown to be increased after exercise. On the other end, there was a reduction in the toxic effects of polyglutamine and amyloid-β peptides after exercise. Exercise was able to positively affect these pathways acutely – after a single hour of thrashing exercise – and chronically, with up to 5 days of exercise in the lifespan studies. Significantly, multiple days of exercise led to benefits in proteostasis that remained days after cessation of exercise. This was observed in both proteasomal function (**Figure 7A**) and the response to amyloid-β (**Figure 9**).

However, the precise mechanism through which exercise modulates amyloid-β in *C. elegans* remains to be determined but may involve a combination of protein chaperones and autophagy. Additionally, the EQ1301 strain appears well-suited to use in the COPAS system for high-throughput screening of drugs, small molecules, and other interventions aimed at increasing autophagy.

These experiments examined different aspects of proteostasis in *C. elegans* following treatment with AKG, a so-called “exerkine,” to see whether it recapitulated the effects of exercise. The divergent effects point to the difficulties in identifying a single “exercise mimetic” or “exercise in a pill”. Part of the difficulty lies in the simultaneity of metabolite changes after exercise: hundreds of metabolites go up and down in different tissues and can travel around the body to be taken up by distant organs – it is possible that the beneficial effects from the elevation of one metabolite only appear in context of a decrease in another. Despite these complexities, these experiments and others show that it may be possible to recapitulate at least partial benefits from exercise through administration of “exercise mimetics”.

A recent study examining the influence of natural genetic variation on polyglutamine toxicity in *C. elegans* introduced a Q40 repeat tagged with a fluorescent cyan protein into a wild isolate from California(64). These wild isolates had a naturally higher level of autophagy due to a variation in the ATG-5 protein. Unexpectedly, higher levels of autophagy were associated with increased aggregation in a tissue-specific manner. They found that by inducing autophagy, they increased aggregation in the muscle (Q40) while reducing aggregation in the intestine (Q40) and neurons (Q67)(64). Comparing this study to my experiments, autophagy was elevated body-wide in *C. elegans* after treatment with AKG and the polyglutamine repeat expressed in these animals is specific to body-wall muscle. This finding of autophagy-dependent aggregation of polyglutamine repeats in the muscle may explain this apparent divergence in proteostasis phenotypes. The mechanisms through which elevated autophagy may both worsen and alleviate aggregation in different tissues remains to be determined, however. Additionally, whether this phenomenon is unique to *C. elegans* or whether this can be found in mammalian systems as well is an open question.

The role of transcriptional and post-translational regulation of autophagy in response to acute exercise is an understudied area. This study found that the TFEB orthologue HLH-30, master transcriptional regulator of autophagy and lysosomal biogenesis, translocates to the nucleus after exercise. Interestingly, this translocation was even higher in food-restricted than exercised animals, although this did not translate to an increase in autophagy in food-restricted animals. The coordination between TFEB and ACSS2 has been recently reported by one study in mice, which found that, in the brain, long-term exercise increased the colocalization and binding of the two proteins in the nucleus post-exercise(65). That study exercised so-called mouse models of Alzheimer’s Disease and found that exercise ameliorated AD brain pathology as well as behavioral deficits. Thus far, this is the only other study to implicate ACSS2/ACS-19 in the response to exercise. Histone acetylation is well-established as a marker of gene activation and increased transcription. Intriguingly, the enzyme ACS-19 (ortholog of ACSS2), which produces acetyl-CoA from acetate, translocated to the nucleus after exercise but not after food-restriction, pointing towards a potential enhancement of transcriptional upregulation via histone acetylation after exercise relative to food-restriction.

## Acknowledgements

Minna Schmidt, Katelyn Adam, Malene Hansen, Lithgow Lab, Andersen lab

